# The Lifespan and Levels of Oxidative Stress between Feral and Managed Honey Bee Colonies

**DOI:** 10.1101/2021.06.29.450441

**Authors:** Kilea Ward, Hongmei Li-Byarlay

## Abstract

Molecular damage caused by oxidative stress may lead to organismal aging and resulted in acute mortality in organisms. Oxidative stress resistance and longevity are closely linked. Honey bees are the most important managed pollinator in agriculture but the long-term survival of honey bees is seriously threatened. Feral honey bee colonies displayed persistence to *Varroa* mites. However, it is unknown whether feral honey bees are stress-resistant or survive longer than managed bee populations. More work is needed to determine the impact of oxidative stress on honey bee health and survival. We used the paired colony design to determine the lifespan and levels of oxidative stress on worker bees from either a feral or a managed colony. Each pair of colonies shared similar foraging resources. Results exhibit longer survival time and lifespans of foragers in feral colonies than the managed colonies. The levels of oxidative stress from the lipid damage of feral colonies are higher than the managed colonies, indicating a tolerant mechanism not a repair mechanism to survive. Our study provided new insights into colony difference of physiology and oxidative stress resistance between feral honey bees and commercial stocks.

## Introduction

Oxidative stress is a redox-sensitive phenomenon that occurs when reactive oxygen species (ROS) are accumulating in a living system faster than their detoxification rate (Farooqui & Farooqui, 2012, Farooqui, 2014, Li-Byarlay & Cleare, 2020). ROS include peroxyl radicals, hydroxyl radicals, hydrogen peroxides, and superoxide anions. ROS levels that exceed the capacity of antioxidant defenses, such as detoxifying enzymes and radical scavenging molecules, cause lipid peroxidation of cell membranes, crosslinking of proteins, DNA fragmentation and damage, and potential cell death (Arking, 2006, Li et al., 2008, Bryden et al., 2013, Johnson et al., 2002, Li-Byarlay & Cleare, 2020).

Pollinators are critical players for the sustainable ecological system because they fertilize a significant number of plants and crops (Kremen et al., 2002, Aizen & Harder, 2009, Burkle et al., 2017, Isaacs et al., 2017, Hunt et al., 2016). The honey bee is the most important managed pollinator for productions of crops and fresh produce (Bond et al., 2014). However, honey bee colony populations are in the steady decline of 30-40% annually, especially in the managed honey bee industry (commercial beekeeping) (Kulhanek et al., 2017). A sustainable beekeeping industry is crucial for agriculture given the increasing demands for pollination as wild pollinator specie populations continue to decline (Aizen & Harder, 2009, Potts et al., 2010). Factors causing the continued decline of honey bee populations are heterogeneous (Neumann & Carreck, 2010) and include pathogens, pesticides, nutrition, genetic variability, and management styles (Nazzi & Le Conte, 2016).

Among and within species substantial variation in ROS susceptibility and longevity exists. However, the mechanisms of this variation have not been elucidated. Specifically, survival advantages during an acute oxidative stress event or normal life history could be conferred by prevention, repair, or tolerance of molecular damage (Li-Byarlay et al., 2016, Li-Byarlay & Cleare, 2020). Feral or wild honey bee colonies live in a non-managed environment and are under natural selection, which persist *Varroa destructor* mites and equipped with higher immunocompetence (Seeley, 2007, López-Uribe et al., 2017, Tarpy et al., 2015). A different environment such as migratory beekeeping management is known to affect the lifespan and oxidative stress in honey bees (Simone-Finstrom et al., 2016, Li-Byarlay & Cleare, 2020). Previous research on oxidative stress and survival in honey bees showed a significant difference of lifespan between honey bee colonies collected in low urbanization environment versus high urbanization environment (Youngsteadt et al., 2015). However, the knowledge of the lifespan, oxidative stress, and aging between feral and managed colonies is unknown. Is it due to different management styles, or different living environments, or different landscape and natural resources? The question addressed by the present study was whether feral colonies display different lifespan or different level of oxidative stress comparing to the managed colonies. The research in survival and oxidative stress will contribute to a valuable explanation of the physiological difference between feral and managed honey bee colonies.

## Materials and Methods

### Bee population and sample collection

A paired colony design was carried out by matching a feral colony with a managed colony within 2 miles of radius (Figure 1). Three paired feral and commercial colonies were sampled in 2018 (Table 1). Feral colonies were in natural habitat of tree cavities. Managed colonies were all originally purchased from commercial package bees in 2018, and maintained by beekeepers with normal beekeeping practice. For each colony, returning foragers were collected by hive entrance by a Heavy-Duty Hand-Held Aspirator (BioQuip, catalog # 2820GA) in mid mornings (10am-12pm), and kept in each insect collecting chamber with end cap (BioQuip, catalog # 2820D) with honey. Live foragers were brought back to the lab and kept in an incubator (model Precision HP GVTY, FisherScientific, Catalog # PR205075G-Q540177) for survival analysis. Additional group of foragers were flash frozen in the field with liquid nitrogen, then kept on dry ice to bring back to the lab and stored in −80 freezer for the analysis of oxidative stress.

**Figure 1.**
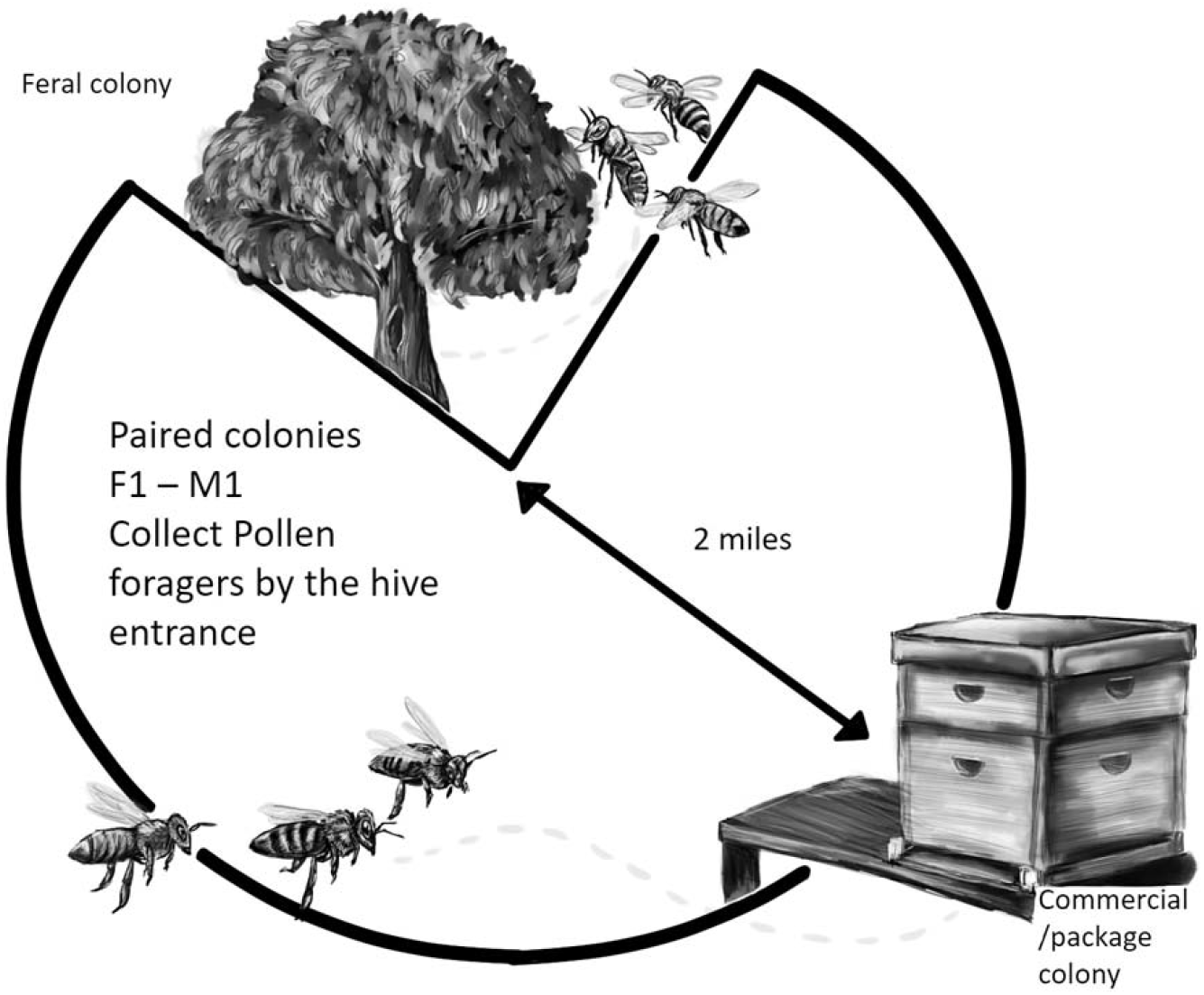
A paired colony design was carried out by matching a feral colony with a managed colony within 2 miles of radius. Managed colonies were maintained by beekeepers with normal beekeeping practice. Feral colonies were in natural habitat of tree cavities.

**Table 1.** Bee colony information from the aging and oxidative stress between feral and managed bees in Ohio.

| Site | Z Stat | P value | Col name | # of feral bees collected | Feral Col GPS Location | # of managed bees collected | Managed Col GPS Location | Date for collection |
| --- | --- | --- | --- | --- | --- | --- | --- | --- |
| Fairborn | 4.28 | <0.001 | F1, M1 | 85 | 39.8412864, -84.0175809 | 50 | 39.8049291, -84.0599239 | Jul 3, 2018 |
| Cedarville | 4.12 | <0.001 | F2, M2 | 174 | 39.7469314, -83.8174640 | 147 | 39.7469314, -83.8174640 | Aug 23, 2018 |
| Bellefontaine | 2.65 | 0.008 | F3, M3 | 106 | 40.395217, -83.620033 | 113 | 40.395217, -83.620033 | July 2, 2018 |

### Experiments for aging and survival analysis

The lifespan of workers from the feral colonies or managed colonies was determined under controlled conditions as described previously (Simone-Finstrom et al., 2016). Lifespan analysis was conducted in August 2018. For each colony type, a group of 20-30 foragers was contained in a 20 oz plastic cup with adequate ventilation in an incubator with 34 degrees and 50-60% of humidity as described before (Evans et al., 2009). A 5 ml syringe was used as a feeder to provide a 50% sucrose solution when needed. The survival analysis of each group showed the lifespan of each colony type (feral or manage).

### Experiments for oxidative stress

ROS-mediated oxidative damage was quantified by measuring MDA level in individual worker heads. The assay was conducted using the OXItek^™^ Thiobarbituric Acid Reactive Substances (TBARS) Assay Kit (ZeptoMetrix Corp). The head of each bee was flash-frozen in liquid nitrogen and immediately pulverized in a 1.5 mL microcentrifuge tube with a sterile plastic pestle. The tissue was then mixed with 280 μ L of PBS, vortexed to homogenize the cellular suspension, and centrifuged briefly to precipitate pieces of tissue and cuticle. Aliquots of the supernatant was used subsequently in the quantification assays. Both the TBARS and the Pierce^™^ BCA^™^ Protein Assay kits (Thermo Scientific) was used according to the manufacturers’ recommendations. Total soluble protein was determined by BCA Protein Assay and used to normalize the corresponding TBARS amounts. Oxidative damage by normalized MDA levels was examined in three biological replicates (colony). Each colony type (feral or package) included at least six individual bees. Total number of samples from feral and managed colonies are 16 and 17.

### Data Analyses

Differences in lifespan between workers from feral and package colonies will be analyzed using the parametric survival analysis JMP Pro 10 with colony treatment (i.e., feral vs. manage) as factors in a general linear model. A log-rank test was used to test for differences in survival. Data of the Oxidative stress was analyzed using a three-way ANOVA to examine if differences in oxidative stress (level of MDA) are due to the population (Feral vs. manage bees). Tukey-Kramer post-hoc tests were used to make pair-wise comparisons of different experimental groups. Differences was considered significant at α = or < 0.05.

## Results

### Survival analysis

To compare colonies, we were able to identify at least three feral colonies in three different locations of Ohio (Table 1). To investigate the difference of lifespan between wild feral colonies and managed package colonies, we performed the survival analysis with live foragers from sampled colonies. For feral colonies, the number of days to survival ranged from 47 to 57 days. In contrast, managed colonies displayed a shorter life span (28 - 42 days) than feral bees (Figure 2). Our data indicated that the lifespan of feral bees was significantly longer than the managed bees (log-rank test; z statistics for pair 1, 2, and 3 are 4.28, 4.12, and 2.65; p values for pair 1, 2, and 3 are < 0.001, p < 0.001, and 0.008) (Table 2).

**Figure 1.**
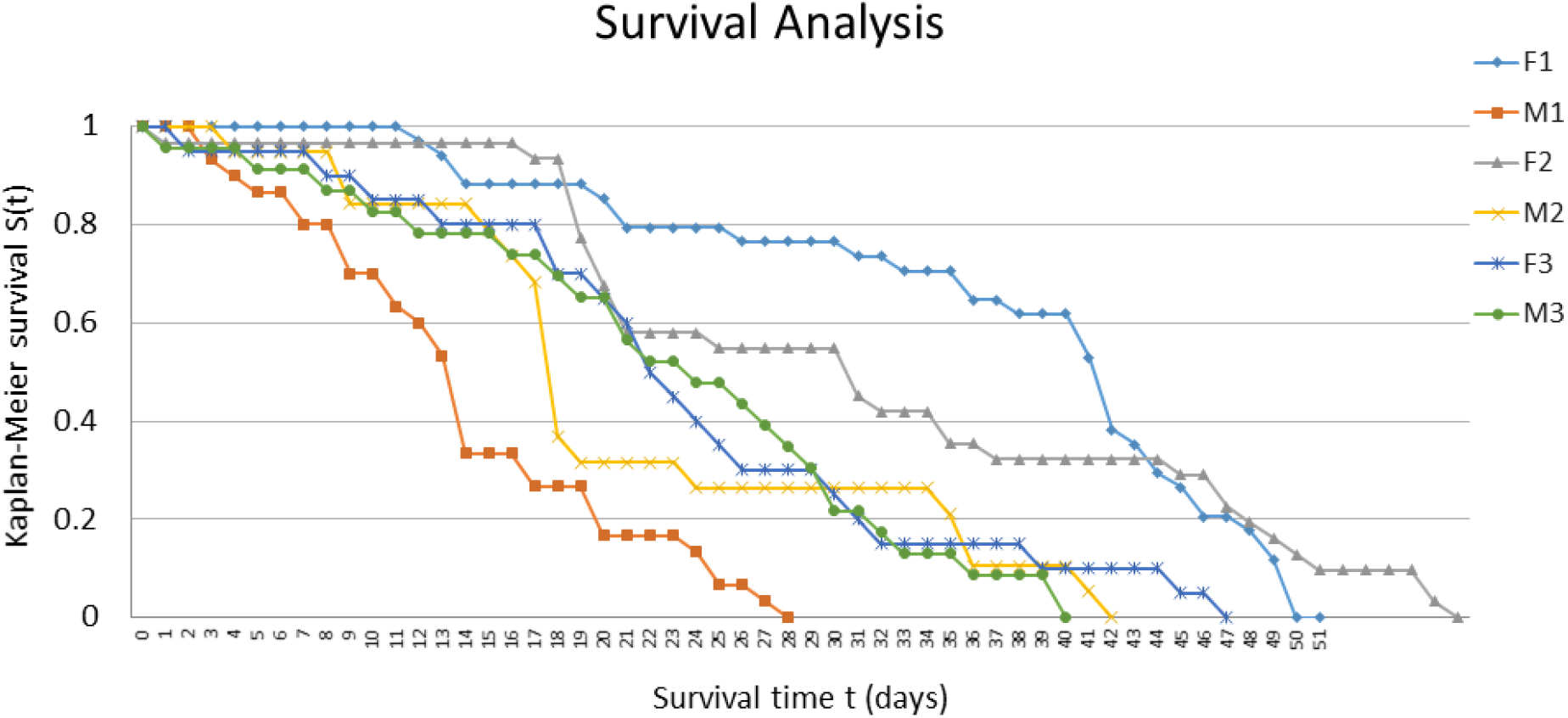
A Kaplan–Meier plot for six groups associated with colony survival, three feral (F1,2,3) and three managed colonies (M1,2,3). Paired colonies (F1-M1, F2-M2, and F3-M3) are within 2 miles. The lifespans of the feral colony were significantly longer than the managed colonies (Kaplan-Meier survival S(t) test, p < 0.001). The x-axis showed the lifespan in days. The y-axis showed the percentage of survival (0-100%). N = 120. Individuals were collected per colony (X2 = 6.48, p < 0.001).

### Lipid damage between feral and managed colonies

To test the level of oxidative stress between two groups, we have collected forager samples from three paired colonies. The oxidative stress in the samples was tested by a TBARS assay measuring the lipid damage as a parameter. The lipid damage indicated by malondialdehyde (MDA) as a major lipid oxidation product was detected using a colorimetric (OD = 532 nm) plate reader. The result in Figure 3 showed a significantly lower level of oxidative stress in feral colonies than package bees (F _1, 32_ = 4.59, p-value = 0.04).

**Figure 2.**
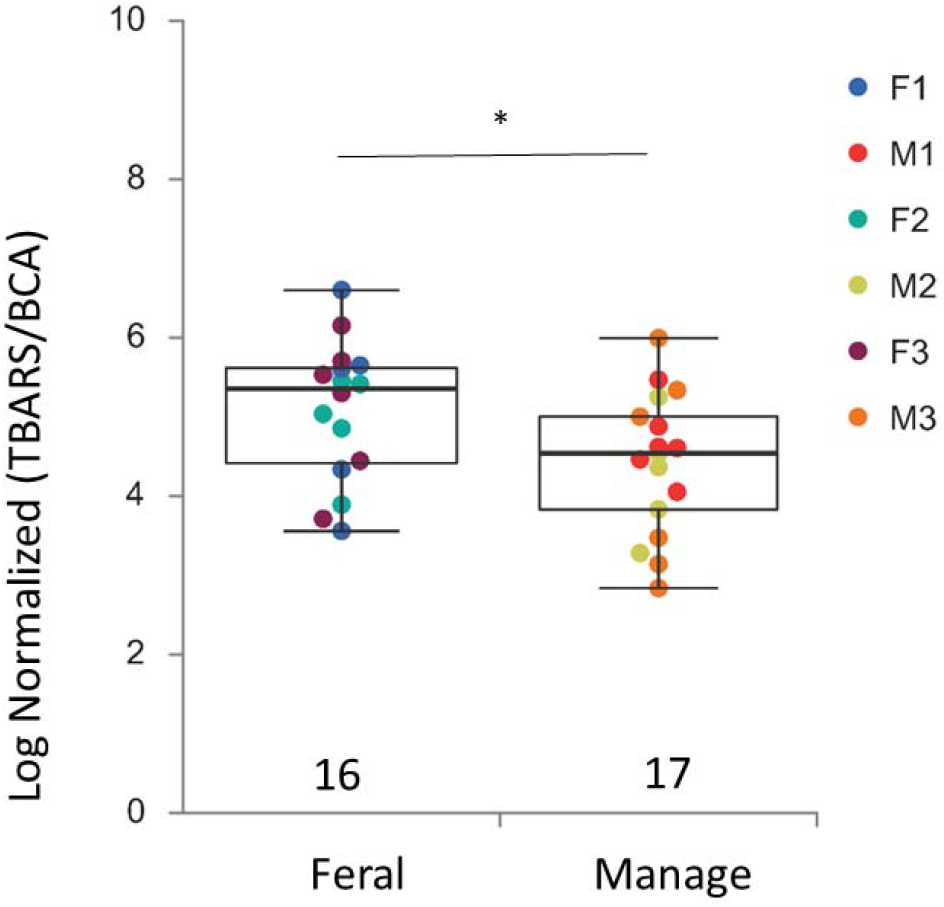
Comparison of an oxidative stress biomarker between feral and managed colonies. Numbers below box-plots are the number of bees tested for each category. p = 0.04.

## Discussion

The aim of this study was to examine the survival and the level of oxidative stress between feral and managed honey bee colonies. The main finding of the study is that the lifespan of foragers from feral honey bees is longer than managed colonies, which is consistent with previous findings in the survival of bee colonies from low urbanization (feral colonies) vs. high urbanization (managed colonies) (Youngsteadt et al., 2015). Feral colonies are wild honey bees and live freely in natural habitats. Also, feral wild colonies provide a resource to study fundamental physiology and ecology of honey bees.

In addition, this study investigated the level of oxidative stress between feral and managed colonies. Our data revealed the level of lipid damage via MDA in feral colonies is higher than managed colonies. This variance indicated different ability between feral and managed honey bees to deal with oxidative stress. A high level of oxidative stress and a long lifespan indicated that feral bees may utilize a tolerant mechanism but not a repair mechanism to deal with oxidative stress. Our results provided the basic knowledge of oxidative stress in wild honey bee population. Additionally, the results support our hypothesis that these oxidative stress levels would be different among different group of honey bee colonies. Only a few studies previously showed different MDA levels were detected between commercial and traditional colonies (Taric et al., 2020) which may be an indication of anthropogenic influence or manipulations. However, there was no comparison of oxidative stress performed between feral and managed colonies. Our data presented here provide the unique information on the lifespan and oxidative stress of feral honey bees.

Previous research indicated that feral honey bee experienced different parasitic pressure than managed honey bees (Thompson et al., 2014), and higher immunocompetency (Appler et al., 2015, López-Uribe et al., 2017). Additional study showed a lower infection rate of trypanosome parasites *Lotmaria passim* (16%) in feral colonies than managed colonies (Williams et al., 2019). These results may also indicate that feral bees experienced different levels of oxidative stress, parasitic, and pathogenic loads from human managed colonies.

Honey bees can show varied responses to oxidative stress depending on their current life stage. Our research only tested forager bees from either feral or managed colonies. Previous study showed that forager brains from summers indicated the highest levels of protein carbonylation as a parameter of oxidative stress and damage than winter bees (Seehuus et al., 2006). It was common to collect foragers to test physiological questions in feral or wild colonies (Appler et al., 2015). Due to the limited access to feral or wild colonies, we were not able to collect nurse bees or young brood for further testing. Foragers display increased oxidative capacity compared to nurse bees because of the energy demands of flight (Harrison, 1986).

Both feralization and domestication can change the selection pressure and population dynamics (Gering et al., 2019). The variation of the phenotypes between feral and managed colonies are due to the natural selection and artificial selection (Parker et al., 2010). For example, previous research showed that feral bee populations that survived *Varroa* mites without miticide-treatment or human interference had lower honey production compared to treated commercial colonies (Le Conte et al., 2007).

It is important to consider that wild colonies may be potential reservoirs for pathogens and mites in adjacent commercial yards (Thompson et al., 2014). Higher levels of deformed wing virus were detected in feral bees compared to commercial (Thompson et al., 2014). *Varroa destructor* genotype traits are also an influence when looking into the oxidative stress and mite pressure on colonies (Anderson & Trueman, 2000). To elaborate, a previous study (Seeley, 2007) found that feral colonies’ survival could have been long lasting due to the mites’ genetic inability to spread harmful viruses instead of the bees’ ability to genetically resist the mites. Locke et al (Locke & Fries, 2011) found no significant difference in brood removal rate, adult grooming rate or mite distribution between feral (unmanaged and surviving with Varroa for ten years) and control managed hives with mite treatments but found the *Varroa* mites themselves had a significant reduction in reproductive success in the feral colonies. This indicated that the natural selective pressure can stabilize the relationship between host-parasite interactions.

